# Rational strain design with minimal phenotype perturbation

**DOI:** 10.1101/2022.11.14.516382

**Authors:** Bharath Narayanan, Daniel Weilandt, Maria Masid, Ljubisa Miskovic, Vassily Hatzimanikatis

## Abstract

Increased availability of multi-omics data has facilitated the characterization of metabolic phenotypes of cellular organisms. However, devising genetic interventions that drive cellular organisms toward the desired phenotype remains challenging in terms of time, cost, and resources. Kinetic models, in particular, hold great potential for accelerating this task since they can simulate the metabolic responses to environmental and genetic perturbations. Although the challenges in building kinetic models have been well-documented, there exists no consensus on how to use these models for strain design in a computationally tractable manner. A straightforward approach that exhaustively simulates and evaluates putative designs would be impractical, considering the intensive computational requirements when targeting multiple enzymes. We address this issue by introducing a framework to efficiently scout the space of designs while respecting the physiological requirements of the cell. The framework employs mixed-integer linear programming and nonlinear simulations with large-scale nonlinear kinetic models to devise genetic interventions in a scalable manner while accounting for the network effects of these perturbations. More importantly, the framework ensures the engineered strain’s robustness by maintaining its phenotype close to that of the reference strain. We use the framework to improve the production of anthranilate, a precursor for pharmaceutical drugs, in *E. coli*. The devised strategies include eight previously experimentally validated targets and also novel designs suitable for experimental implementation. As an essential part of the future design-build-test-learn cycles, we anticipate that this novel framework will enable high throughput designs and accelerated turnover in biotechnological processes.

## Introduction

Advances in gene editing techniques and the ever-increasing availability of omics data have spawned intense efforts in metabolism research. Within the biomedical domain, this has enabled us to glean broader insights into the metabolic phenotypes of various diseases, allowing for more informed therapeutic interventions^1–3^. In biotechnology, these advances have led to the creation of environmentally friendly, cost-effective bio-foundries using genetically engineered cellular organisms for optimal production of valuable compounds^4^. These metabolic engineering undertakings are typically implemented as a design-build-test-learn cycle, involving multiple experimentation stages and fine-tuning strain designs^5^.

While technological advances have facilitated the genetic manipulation of organisms, significant challenges remain in determining the targets, and the extent, of such manipulations. Since robustness to changing environmental conditions is essential for the viability of designed strains, we need to ensure that the genetic interventions maintain critical cell properties such as the energy charge and redox potentials^6–9^. To achieve this, it is typically necessary to develop strategies targeting more than one reference strain enzyme. Unfortunately, devising such multi-target strategies by direct experimentation requires considerable time and resources. One approach to reducing these costs is to conduct rational metabolic engineering using computational models to narrow down the range of strategies to be experimentally verified.

In particular, dynamic metabolic models are well suited for this task since they can capture the temporal evolution of the metabolic states to environmental and genetic perturbations under real-world fermentation conditions. However, the lack of available information about the values of kinetic parameters hampers the development of these models. Indeed, even for well-studied organisms such as *E. coli* or *S. cerevisiae*, we can find experimentally obtained values for only a few parameters in the literature and databases^10,11^. To infer the values of missing kinetic parameters, researchers have traditionally employed parameter estimation^12–14^ and Monte Carlo techniques^15–19^. Recently, there have been numerous efforts to use machine learning to accelerate the building of these models^20–23^.

Even when a high-quality kinetic model is available, it is computationally challenging to determine targets for metabolic engineering that meet desired design specifications because it requires simulating the metabolic network’s responses for many putative designs. For example, one would need to perform more than 2.5 · 10^9^ simulations to exhaustively explore all possible strategies for manipulating five enzyme activities within a middle-sized metabolic network of 200 reactions (catalyzed by 200 enzymes). Additionally, for all these simulations, we would need to analyze whether or not the designed strains meet the specifications and preserve the robustness of wild-type strains exposed to long-term evolutionary pressure^24^. Hence, to perform reliable and comprehensive strain designs, the research community needs systematic, resource-efficient approaches that leverage the predictive capabilities of nonlinear kinetic models.

In this work, we report NOMAD (NOnlinear dynamic Model Assisted rational metabolic engineering Design), a computational framework that scouts the space of candidate metabolic engineering strategies and then proposes those that satisfy the desired design specifications while maintaining the robustness of the engineered strain. NOMAD preserves the robustness of the engineered strains by maintaining their physiology close to the original phenotype shaped through the course of evolutionary pressure and selection. As it has been hypothesized and shown earlier^25,26^, we also found here that this can be achieved by maintaining their metabolite concentrations and fluxes close to those of the reference strain. The rationale of trying to ensure a minimal deviation of the engineered strain phenotype from that of the reference strain has also been put forth in a constraint-based modeling approach called MOMA^24^. The departure in this work is that kinetic models couple metabolite concentrations, metabolic fluxes, and enzyme levels, thus allowing us to represent the studied phenotype with higher fidelity and capture both steady-state and dynamic properties of the studied metabolic phenotype. Additionally, NOMAD proposes testing the sensitivity and performance of the designs in nonlinear dynamic bioreactor simulations that mimic real-world experimental conditions. We can then suggest the best performers from these tests for experimental validation with high confidence. As a validation study, we use this method to propose engineering targets for the overproduction of anthranilate in a previously studied strain of E.coli^27^. NOMAD proposed four candidate designs that proved robust across phenotypic and expression uncertainty while also providing a superior in silico performance when compared with experimentally devised strategies^27^.

Through its conception, NOMAD lends itself well to the DBTL (design – build – test – learn) cycle, with every round of iteration improving the quality of the proposed strain designs. Overall, NOMAD has the potential to accelerate the pace at which strain design breakthroughs are achieved, representing a potent disruptor within the biomedical and biotechnological domains.

## Results

### NOMAD for reliable strain designs

The NOMAD workflow consists of three steps (Figure 1). Briefly, it starts by generating a population of putative kinetic models consistent with experimentally observed omics and cultivation data, physicochemical laws, network topology, and regulatory interactions. These kinetic models consist of a system of nonlinear ordinary differential equations (ODEs) characterized by a set of kinetic parameters. To generate such models, we can use traditional kinetic modeling approaches such as MASSPy^17^, Ensemble Modeling^19^, ORACLE^15,18,28,29^, and machine learning empowered methods such as REKINDLE^21^ and iSCHRUNK^20,22^.

**Figure 1:**
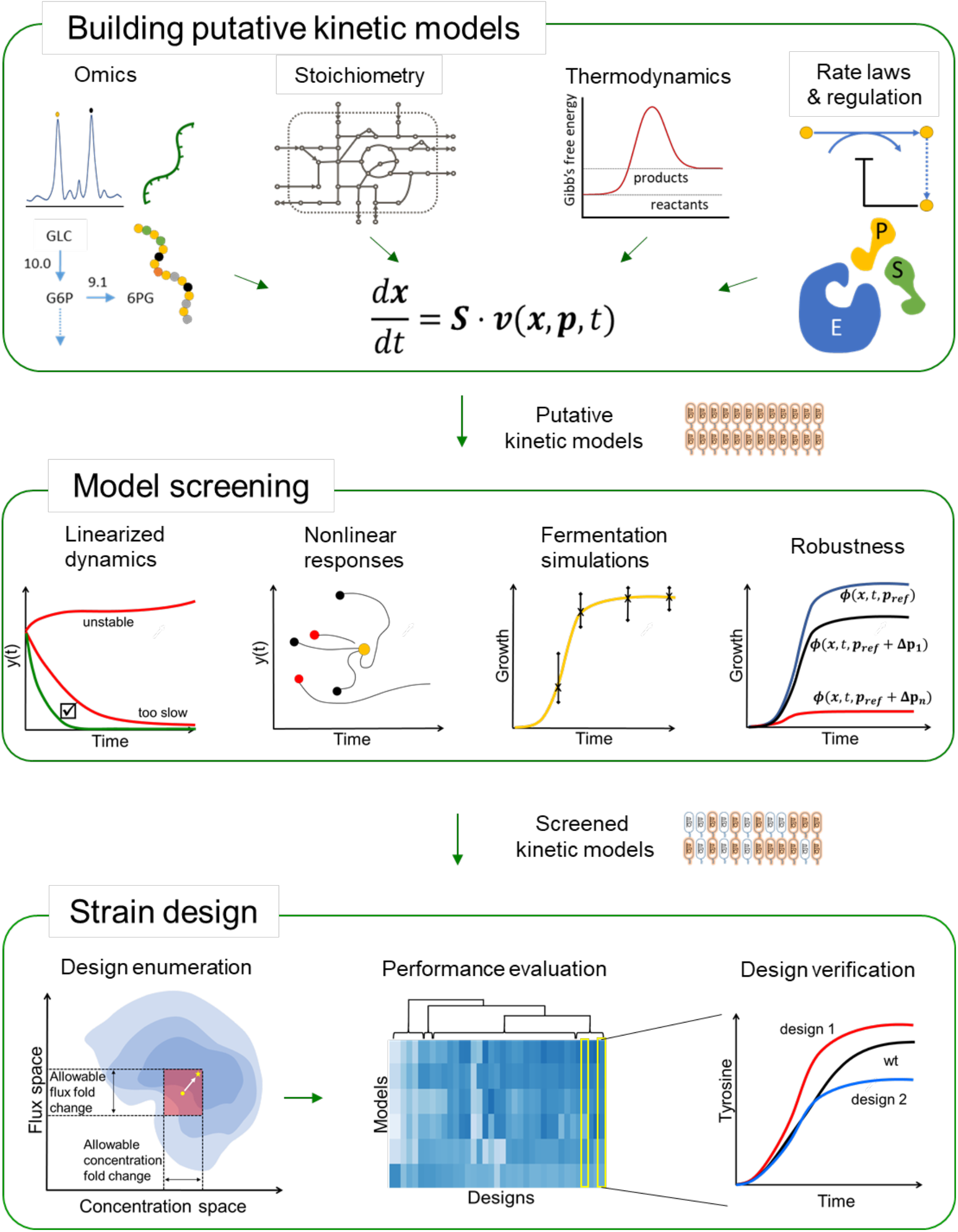
Steps of the NOMAD workflow. We first build a set of putative kinetic models, represented by a system of ODEs, by integrating different types of data. Next, we choose models based on dynamic characteristics such as their stability, ability to reproduce experimental fermentation data, and robustness to enzymatic interventions. In the final step, we use the chosen models to conduct strain design. This involves solving a MILP optimization problem and enumerating designs that maintain the engineered strains close to the reference strain, evaluating the performance of these designs, and verifying the top designs in silico before sending them to experimentalists.

In the second step of NOMAD, we perform several quality checks on the kinetic models and identify those that will ensure reliable in silico strain design strategies^30^. In this screening process, we retain kinetic models that are (i) consistent with experimentally observed steadystate values of metabolic fluxes and metabolite concentrations, (ii) locally stable around that steady state; (iii) able to reproduce the dynamic behavior of metabolic responses under industrial production conditions; (iv) consistent with any available information on studied phenotype or a piece of expert knowledge; and (v) robust, meaning that these models resist change and are capable of coping with various genetic and environmental perturbations.

In the final step, we use the screened models to design engineering strategies for achieving a chosen metabolic objective, such as the overproduction of high-value biochemicals. We use Network Response Analysis^26^ (NRA) to perform the strain design. NRA casts the strain design process as an optimization problem that uses the outputs of the kinetic models (Methods) and integrates design constraints ranging from the allowable fold changes in concentrations and fluxes to the extent and number of enzymatic interventions. This way, we obtain a computationally efficient modus operando to enumerate designs and maintain the physiology of the engineered strain close to the reference physiology through various constraints.

After enumerating alternate strategies, we test their performance and sensitivity to phenotype and expression variability in fermentation simulations that mimic real-life conditions. We finally propose the best-performing designs to experimental partners for implementation.

NOMAD is described in more detail in the Methods section.

### Improving anthranilate production in *E. coli*

As a case study for testing and validating the NOMAD workflow, we designed strategies for increasing anthranilate production in *E. coli* strain *W3110 trpD9223*. In an earlier experimental study, the strain was used as a scaffold for overproducing anthranilate through several genetic manipulations^27^.

#### Kinetic models representative of *E. coli W3110 trpD9923*

We used ORACLE^15,16,28,31^, implemented in the SKiMPy toolbox^32^, to generate a population of 800,000 putative kinetic models that satisfied the experimentally observed steady-state behavior of the reference strain (Methods). However, not all of these models were necessarily suitable for strain design due to poor dynamic characteristics or poor responses to engineering interventions, necessitating the process of model screening. We first screened the population of kinetic models for those with dominant time constants, quantified by the inverse eigenvalues at the steady state, at least 5x faster than the doubling time of the cell (Methods). This meant that all metabolic processes were *expected* to settle into their steady states within the doubling time of the cell (Methods). More than 11% of the generated models (91,852) showed such dynamic characteristics.

Whereas the inverse eigenvalues are a good indicator of the dynamic of metabolic responses in close vicinity of the steady state, due to the nonlinear nature of the system, not all models will exhibit the same dynamic behavior in a batch setting where metabolic states change intensely. Therefore, we tested if these models could reproduce experimentally observed behavior in a batch reactor. Out of 91,852 models, 212 captured experimentally observed temporal evolutions of growth, anthranilate, and glucose (Figure 2B–D).

**Figure 2:**
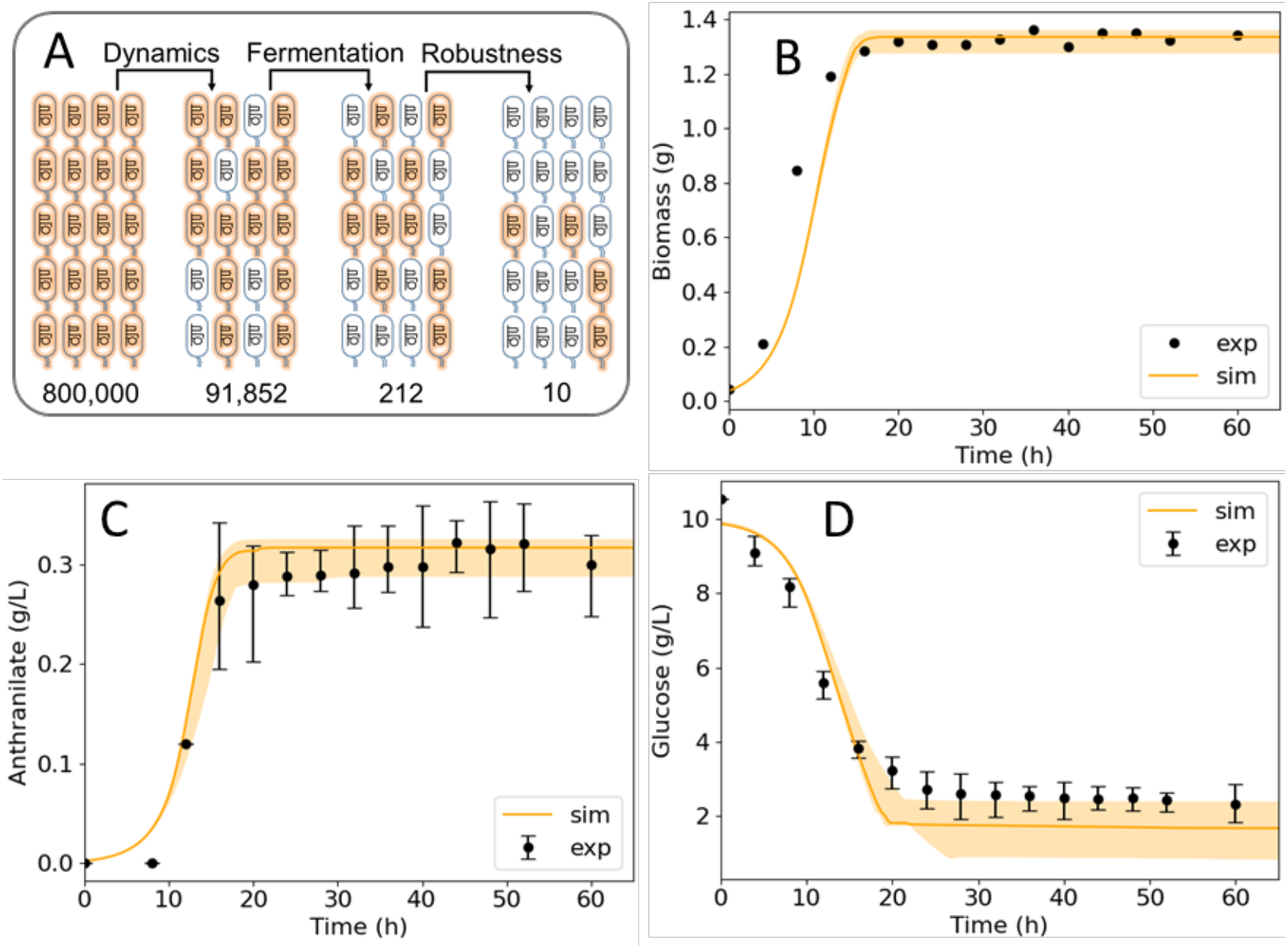
Results of the model screening process. (A) We gradually filter the models from 800,000 putative kinetic models to 212 that could reproduce experimentally observed fermentation curves. 10 of these 212 models also proved to be robust to enzymatic perturbations. The median (solid line) and spread (shaded region) of the growth (B), anthranilate (C) and glucose (D) time evolution of these 10 models (solid line) show close match with experimentally obtained data.

Next, we evaluated the suitability of these 212 models for strain design by testing their responses to naturally occurring random perturbations in enzyme activities. 10 out of the 212 models proved to be robust and consistent with the studied strain, exhibiting at least 50% of the growth of the reference strain (Methods) when subjected to these perturbations. The responses of these 10 models in a batch fermentation setup show qualitative and quantitative agreement with the experimentally observed time evolutions of growth, anthranilate production, and glucose consumption (Figure 2).

#### Closeness to the reference physiology is essential for the performance of designed strains

Extensive metabolic engineering might steer the engineered strain towards metabolic states with impeded growth or performance as, too often, the objective of overproduction of a metabolite has a major tradeoff with the organism growth and global biosynthetic processes. For example, in efforts to optimize the specific productivity or yield of target chemicals, metabolic engineering interventions could inadvertently reduce other cell capabilities such as ATP production or redox potential by redirecting carbon and biosynthetic resources towards the target pathways and products. Because reference strains have evolved to maintain healthy and robust physiology, we postulate that we can engineer a productive strain by redirecting the flux to the desired objective while remaining as close to the reference state as possible. Thus, we use the proximity of the metabolic and fluxomic profile of the engineered strain as a proxy for maintaining a vigorous phenotype^25,26,33^. A related concept was also studied in the context of steady-state flux analysis.^24^ Here, NOMAD uses nonlinear kinetic models and network response analysis (NRA, see Methods) to implement this concept and constrain the phenotype perturbation while maximizing productivity.

To verify the postulated hypothesis, we used NRA for the 10 chosen kinetic models to design several groups of strains with an improved yield of anthranilate on glucose. The groups differed by how much the engineered strains could deviate from the reference strain, quantified through a fold-deviation of the intracellular metabolite concentrations and metabolic fluxes from their values at the reference state. The group of strains closest to the reference strain could have the intracellular concentrations deviating 2-fold from the reference strain. In contrast, the less constrained group could have the intracellular concentrations deviating 20-fold from the reference strain. For all the groups, we allowed up to three enzyme modifications with a maximum of 5-fold change in their activities.

Current approaches to strain design using kinetic modeling perform a metabolic control analysis (MCA)^34,35^ around the reference state and rank the target enzymes using the absolute value of the product flux or yield control coefficient with respect to each enzyme in the network. This approach does not take into account constraints that could maintain the healthy physiology we discussed above. To understand the implications of using unconstrained MCA for strain design, we also applied a 5-fold change in enzyme activities to the enzymes corresponding to the top 3 anthranilate yield control coefficients for each kinetic model, without any constraints on concentration and flux perturbations.

The nonlinear responses of all these engineered strains showed that the closeness to the reference strain impacted performance. Indeed, for the groups closest to the reference phenotype (2-fold and 3-fold deviations), the engineered strains retained the dynamic characteristics of the reference strain while producing higher anthranilate titers (>15%) at a modest cost to growth (<16%). Moreover, the titers achieved by these strains (~0.38 g/L) are only attained by the designs with 10-fold and 20-fold deviations after more than double (~40 hours) their fermentation time.

In contrast, the designs with 10-fold and 20-fold deviations demonstrated slower dynamics than the reference strain, with lower mean titers and growth at the end of the production period for the latter (Figure 3). In a similar vein, the unconstrained designs (‘MCA’ in Figure 3) consistently failed to achieve any semblance of growth or production of anthranilate. The likely explanation for this is that, by not constraining the phenotype perturbation, we pushed the engineered strains far away from the reference strain while disregarding the network effects of the enzyme modifications. Even when we considered the targets that had a nonnegative impact on growth, the resulting responses were inferior to the designs generated using NRA (Supplementary note V).

**Figure 3:**
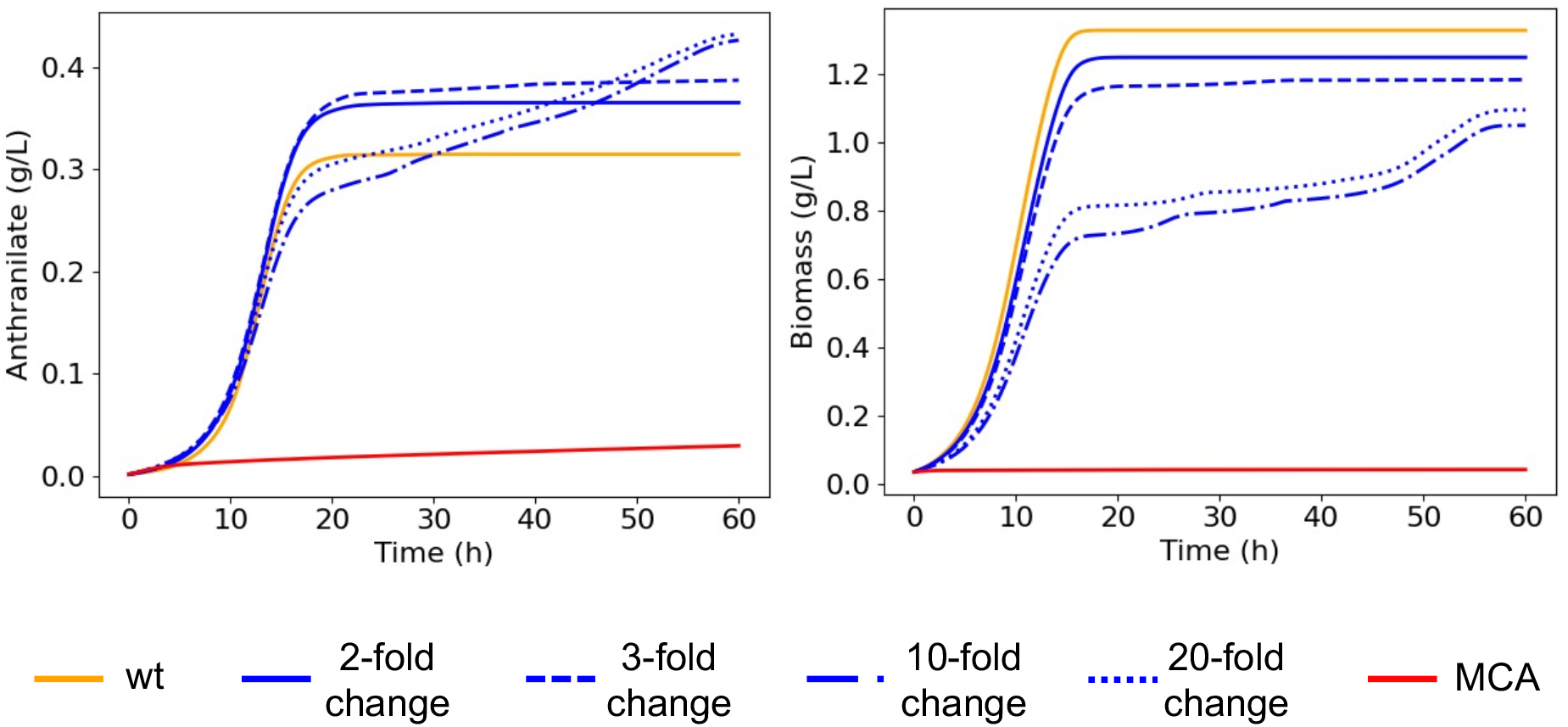
Results of the phenotype perturbation study showing mean anthranilate and biomass responses of engineered strains under different allowable fold changes in concentrations with respect to the reference strain (wt). As we permit a greater deviation from the reference physiology, from 2-fold (solid blue) to 20-fold (dotted blue) changes in concentrations, we observe a decrease in the titers of anthranilate and biomass across the 10 models at the end of the fermentation period of the reference strain (18 hours). Furthermore, the completely unconstrained approach to strain design that uses the top-3 control coefficients alone (MCA) stifles growth as well as anthranilate production. These results underline the importance of adhering to the reference physiology when conducting strain design.

These results suggest that for the cost-effective production of valuable biochemicals, it is judicious to generate designs that minimize phenotype perturbation while respecting other design specifications such as maximal titer and specific productivity.

#### NOMAD designs encompass experimentally validated targets around shikimate metabolism

We employed NRA^26^ to engineer strains with a maximized yield of anthranilate with respect to glucose uptake for each of the 10 kinetic models (Methods). Using the results of the phenotype perturbation study, we permitted no more than a 3-fold change in concentrations to ensure we adhered to the phenotype of the reference strain. In addition, we allowed three enzyme modifications with a maximum 5-fold upregulation, and unrestricted downregulation. There were multiple designs for each model within 5% of the maximal anthranilate yield. The number of such design alternatives for each model ranged from 2 – 12. In total, we obtained 70 designs involving 37 enzymes across the 10 models predicting a 90-158% increase in anthranilate yield. Out of the 70 designs, 41 were unique by membership, meaning that they contained a unique set of three enzymes to be targeted (Figure 4).

**Figure 4:**
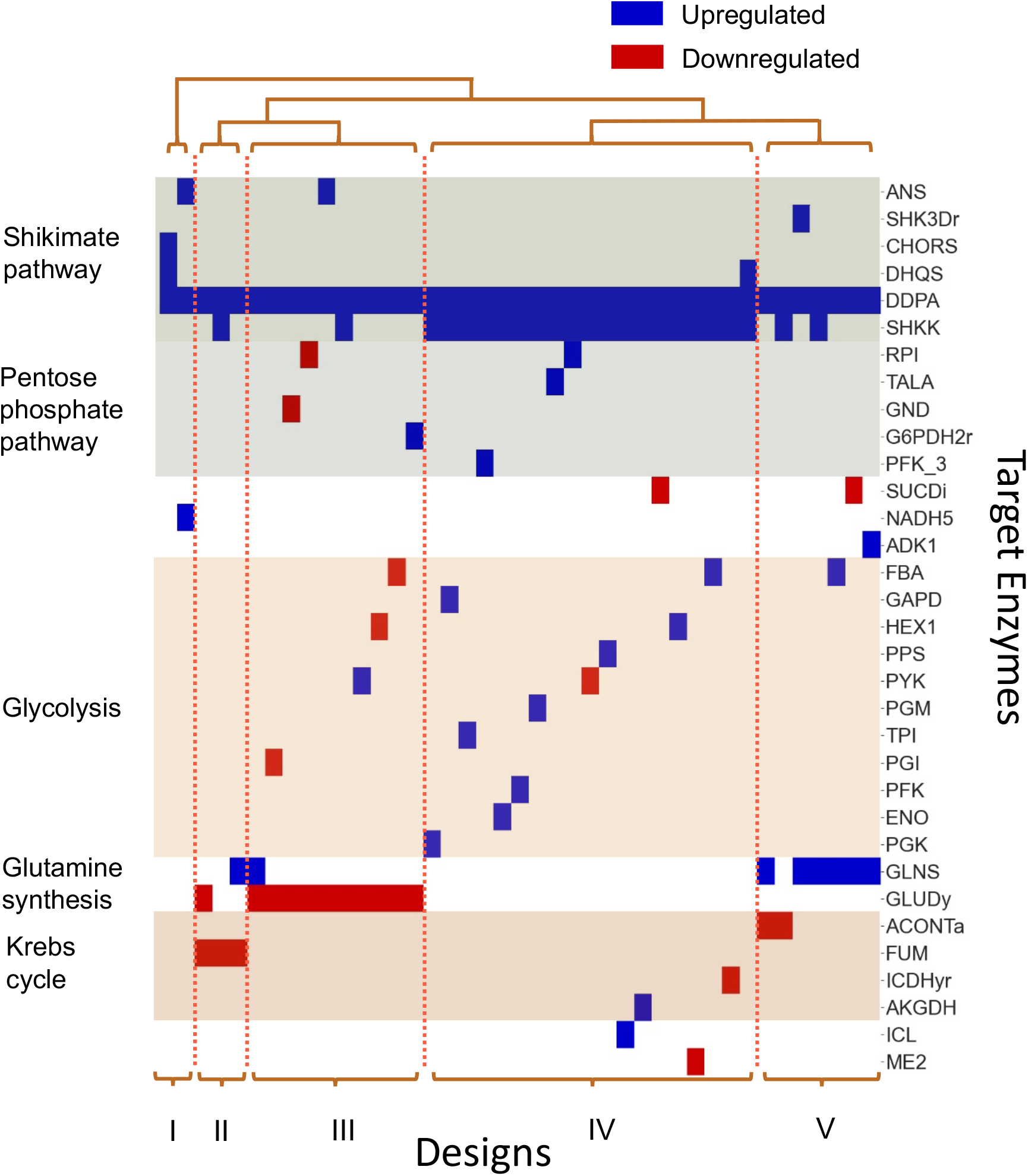
Clustering of the 41 unique NRA designs. Each design contains a unique set of enzymes to be targeted. This clustering analysis revealed 5 distinct manners for overproducing anthranilate. **Abbreviations:** SHKK: Shikimate kinase, CHORS: Chorismate synthase, DHQS: 3-dehydroquinate synthase, CHORM: Chorismate mutase, ANS: Anthranilate synthetase, ANPRT: Anthranilate phosphoribosyltransferase, DDPA: 3-deoxy-D-arabino-heptulosonate 7-phosphate synthetase, GND: Phosphogluconate dehydrogenase, PFK_3: Phosphofructokinase (s7p), RPI: Ribose-5-phosphate isomerase, G6PDH2r: Glucose 6-phosphate dehydrogenase, NADH5: NADH dehydrogenase, SUCDi: Succinate dehydrogenase (irreversible), HEX1: Hexokinase, PGK: Phosphoglycerate kinase, PYK: Pyruvate kinase, PGM: Phosphoglycerate mutase, PGI: Glucose-6-phosphate isomerase, PPS: Phosphoenolpyruvate synthase, GAPD: Glyceraldehyde-3-phosphate dehydrogenase, FBA: Fructose-bisphosphate aldolase, PFK: Phophofructokinase, TPI: Triose-phosphate isomerase, FBP: Fructosebisphosphatase, ENO: Enolase, PSERT: Phosphoserine transaminase, GLUDy: Glutamate dehydrogenase, GLNS: Glutamine synthetase, ACONTa: Aconitase, AKGDH: 2-Oxogluterate

Remarkably, 8 out of the 37 enzymes involved in the designs were validated experimental targets for increasing the flux through the shikimate pathway^36–38^. Three out of the 8 enzymes, DDPA, DHQS, and SHKK, belonged to the shikimate pathway, three belonged to glycolysis, PGI, PPS, and PYK, and two belonged to the pentose phosphate pathway, TALA and G6PDH2r (Figure 4). More specifically, Patnaik et al. increased the carbon flow through the shikimate pathway by overexpressing DDPA (aroG) alone and DDPA along with PPS (ppsa)^38^. All our designs target DDPA, and one in particular targets both DDPA and PPS, along with SHKK (aroK). Rodriguez et al. reviewed strategies that sought to increase the production of aromatic amino acids by either increasing the availability of the precursors to the shikimate pathway or enhancing the activity within the pathway^37^. The reviewed strategies directly targeting the shikimate pathway included the deregulation of DDPA, DHQS (aroB), or SHKK. Among the experimental strategies that targeted phosphoenolpyruvate (pep) or erythrose-4-phosphate (e4p) availability were those that inactivated the pyruvate kinases (pykAF), increased the activity of PPS, knocked out PGI (pgi), or redirected carbon to the pentose phosphate pathway (PPP) through the upregulation of TALA (talB), TKT (tktA), or G6PDH2r (zwf). NOMAD designs contained all previously mentioned interventions, except for TKT.

Interestingly, although one of the generated designs proposes PYK downregulation in line with the experimental approach (Figure 4, DDPA↑, SHKK↑, PYK↓), another design proposes its upregulation instead (DDPA↑, PYK↑, GLUDy↓), suggesting the possibility of alternative regulation patterns when targeting multiple enzymes simultaneously.

In addition to encompassing several reported experimental interventions, NOMAD also suggested novel targets that can achieve the same impact on anthranilate production as the expert-proposed candidates. Some of them frequently appeared in our designs, such as the downregulation of GLUDy (11/41 designs) and the upregulation of GLNS (8/41 designs). In contrast, the upregulation of ENO and ICL appeared only in one design each.

#### NOMAD performs well compared to experimental strategies for anthranilate production

To evaluate the performance of NOMAD designs against those reported experimentally, we implemented two of the engineered strains reported by Balderas-Hernandez et al.^27^ in a bioreactor setup. For the experimental implementation of the first strain, W3110 trpD9923/pJLaroGfbr, the authors used a feedback resistant version of *aroG* to redirect carbon into the Shikimate pathway, resulting in an increase in anthranilate titers from 0.31g/L to 0.4g/L. For the second strain, W3110 trpD9923/pJLaroGfbrtktA, they additionally increased the availability of e4p through the overexpression of transketolase (associated with TKT1 and TKT2), resulting in titers of 0.7g/L.

We used the 10 kinetic models to simulate the feedback-resistant version of aroG by removing the inhibition of DDPA by phenylalanine. For transketolase overexpression, we applied a 5-fold increase in the enzyme activities of TKT1 and TKT2. Although our models provided lower median titers of anthranilate than those reported experimentally, they captured the performance trends of both interventions – trpD9923/pJLaroG^fbr^tktA produced a better median titer (0.36g/L) than trpD9923/pJLaroG^fbr^ (0.33g/L) which was in turn superior to trpD9923 (0.31g/L) (Figure 5). The obtained time evolutions for the glucose uptake and growth reached their final values faster than the experimental observations (Supplementary note IV). Interestingly, although we did not integrate information about the two engineered strains in the model-building process, our models could reproduce the experimental observation that the difference in anthranilate titers between the two engineered strains was greater than the difference between the wild-type and trpD9923/pJLaroG^fbr^.

**Figure 5:**
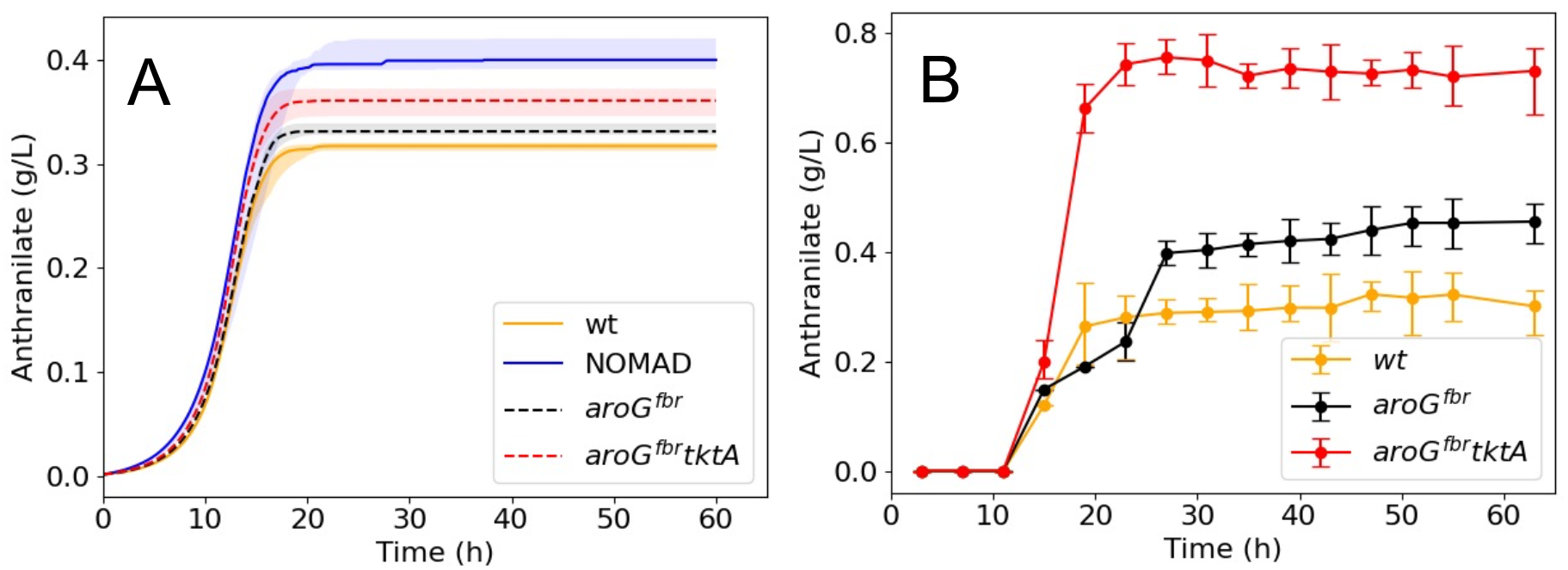
A comparison of simulated (A) vs experimentally observed (B) responses for W3110 trpD9923 (orange), W3110 trpD9923/pJLBaroG^fbr^(black) and W3110 trpD9923/pJLBaroG^fbr^ tktA (red). The median (solid) and 1^st^ and 3^rd^ quartiles (shaded) of the responses over the 10 kinetic models captures the trends reported experimentally, with the overexpression of tktA resulting in a superior titer of anthranilate when compared to the targeting of aroG alone. With the NOMAD designs (blue) we obtain a superior titer of anthranilate when compared with the in silico implementation of the experimental designs.

To benchmark the engineering strategies suggested by NOMAD, we also compared the simulated responses of the experimental strains to the top 5 designs applied using the NRA suggested fold changes specific to each model. We found that the NOMAD designs resulted in superior median anthranilate titers (0.4g/L) compared to the simulated experimental approaches, suggesting that the strains W3110 trpD9923/pJLaroG^fbr^ and W3110 trpD9923/pJLaroG^fbr^tktA could be further improved (Figure 5).

#### Alternative routes for producing anthranilate

To detect common patterns and routes toward producing anthranilate across the designs, we conducted a clustering analysis of the 41 unique designs (Figure 4). This revealed the presence of five clusters of alternative enzymatic interventions satisfying the imposed design specifications. All the designs redirect carbon to the shikimate pathway by increasing the activity of DDPA which serves as the entry point to the pathway. The clusters differed by the choice of the other two target enzymes. Cluster I consists of two designs, one of which concentrates the flow of carbon through the shikimate pathway (CHORS and DHQS), while the other increases the activity of anthranilate synthase (ANS) and the activity of NADH5 in the electron transport chain (ETC). Cluster II has three designs, all of which reduce the activity in the Krebs cycle (FUM). Two of the designs also increase the availability of glutamine, which is a substrate for ANS, either by increasing its synthesis (GLNS) or decreasing the conversion of glutamate to *α*-ketoglutarate (GLUDy) so that it is available for glutamine synthesis. The third design increases the activity of SHKK in the shikimate pathway.

Cluster III contains designs that all target the availability of glutamate for glutamine synthesis by reducing its degradation (GLUDy). Some of the designs also balance the availability of the precursor metabolites, e4p and pep, by targeting enzymes in the PPP or glycolysis. The others target either the activity in the shikimate pathway (SHKK, ANS) or the availability of glutamine (GLNS).

The largest cluster, cluster IV, has designs that focus on the shikimate pathway by increasing the activity of SHKK. Additionally, in a manner similar to cluster III, some designs in this group ensure the balance between the two shikimate pathway precursors by targeting glycolysis (ENO, FBA, etc.) or PPP (TALA, RPI). The remaining designs target either growth, through the ETC (SUCDi), the Krebs cycle (ICDHyr, AKGDH), or anaplerotic reactions (ME2, ICL), or the production of anthranilate through the shikimate pathway (DHQS) (Figure 4).

Finally, cluster V is an agglomeration of designs that focus on glutamine synthesis with all but one of the designs targeting GLNS. Additionally, the designs target enzymes in the shikimate pathway (SHKK, SHK3Dr), the ETC chain (SUCDi, ADK1), the Krebs cycle (ACONTa) and glycolysis (FBA).

NOMAD identified a set of multiple alternative designs that would improve anthranilate production to a similar extent. In the next section, we showcase a procedure for identifying this set’s most robust and implementation-suitable designs. Further expert knowledge can be used to perform a comparative analysis of the alternative routes and select which designs to implement experimentally.

#### Robust and implementation-suitable designs

As we cannot know, a priori, which among the population of kinetic models best represents the physiology of the cell, it is judicious to ensure that the proposed engineering strategies are reliable and consistent across the range of phenotypes spanned by the 10 kinetic models. To this end, we implemented the 41 unique designs (Methods) in each kinetic model and evaluated the mean increase in anthranilate yield predicted by NRA across all the models. The five designs with the highest predicted mean increase (~93%) all belonged to Cluster III (Figure 6A). They all suggest redirecting carbon to the shikimate pathway by upregulating DDPA, and increasing the availability of glutamate for glutamine synthesis by downregulating GLUDy (Figure 6B). Four of the designs also balance the availability of pep and e4p by targeting glycolysis (HEX1, PYK, PGI) or the pentose phosphate pathway (GND). The fifth design increases the enzyme activity of ANS which is responsible for anthranilate synthesis.

**Figure 6:**
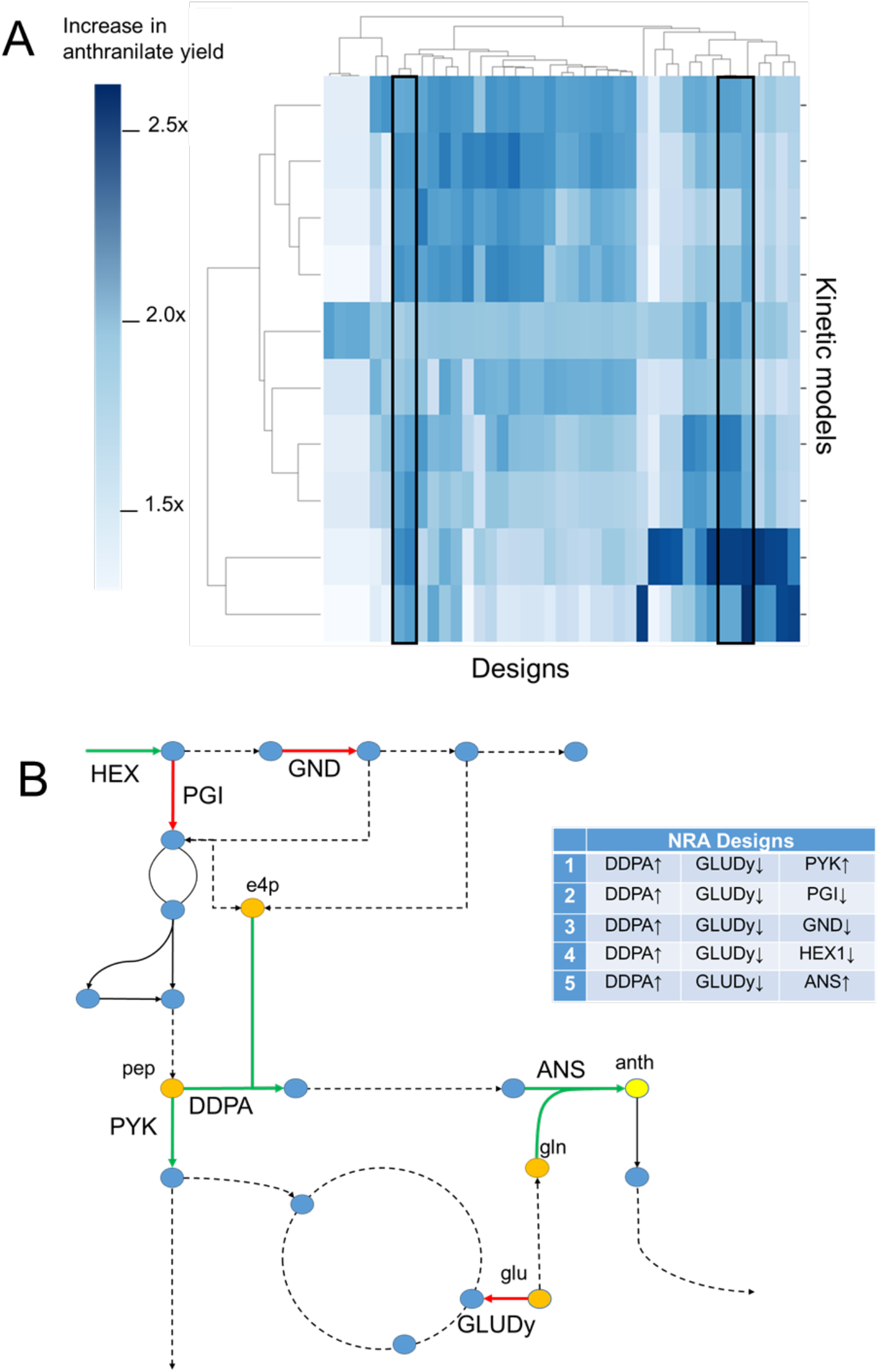
Design evaluation and pruning. (A) A heat map of the predicted increase in anthranilate yield when each of the 41 designs (columns) is applied to the 10 kinetic models (rows). The five designs with the highest mean NRA solution are marked in black rectangular frames. (B) A schematic of the metabolic network containing the target enzymes from these five designs. The designs target the activity of DDPA and GLUDy, while differing by the third target enzyme.

Before recommending any design for experimental implementation, we must test and validate them in nonlinear simulations that closely mimic real-world conditions (Methods). Hence, we applied these designs to all 10 models in a batch fermentation setting, using the fold changes suggested by NRA for each combination of model and design (Figure 7A and 7B). Four out of the five designs (d-1 – d-4) performed well across the phenotypic uncertainty covered by the 10 models. They remained close to the phenotype of the reference strain while providing >25% increases in anthranilate titers, as shown by the mean of their responses across the 10 models (Figure 7). Design d-5 (DDPA, GLUDy, ANS) was discarded due to its poor performance across the models - it displayed significantly slower dynamics and only reached the anthranilate titers of the reference strain after 40 hours.

**Figure 7:**
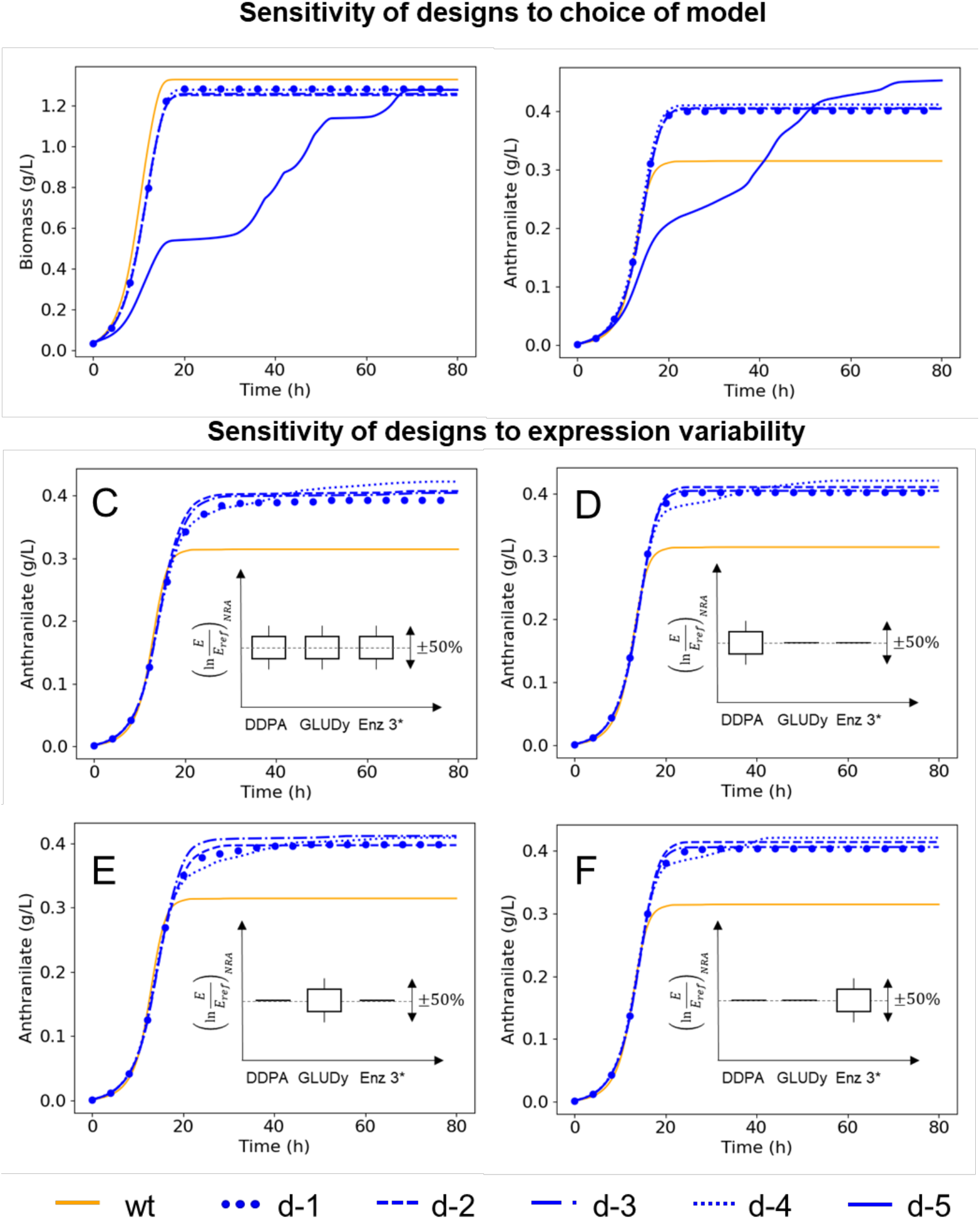
Validation and sensitivity analysis of the top five designs in a fermentation setting. The mean responses across the 10 kinetic models when the five designs are applied using the NRA suggested fold changes specific to each model (A and B). Four out of the five designs retained their performance across the models, with only d-5, targeting DDPA, GLUDy, and ANS, demonstrating significantly altered dynamics. These four designs also proved to be robust to errors in experimental implementation, maintaining their performance when subjected to a ±50% perturbation applied to their mean suggested NRA fold changes for all the enzymes (C), and for each enzyme individually while keeping the other fold changes at the mean NRA suggested values (D, E, F).

The outcome of the experimental implementation of these designs while in agreement with the predicted trends, it will most likely deviate from the NRA-predicted responses to the corresponding fold changes in enzyme activities. Hence, in order to quantify the expected deviations, we conducted a global sensitivity analysis by applying perturbations to the mean NRA-proposed fold changes in enzyme activity (Supplementary note II), and we computed the margin for error afforded by each of the four robust designs. We found that all these designs could withstand errors in experimental implementation, retaining their performance even when we applied ±50% perturbation to all three enzymes together (Figure 7C and Supplementary note III) and to each enzyme individually while keeping the others at the mean NRA-suggested values (Figures 7D–F and Supplementary note III). By retaining their performance across a range of models, and a spread of enzyme expression levels, the four screened designs, DDPA + GLUDy + PGI/GND/HEX1/PYK, proved to be robust to physiological and expression level uncertainties and can thus be confidently passed on for experimental validation.

The results of the *in silico* validation studies underline one of the key features of NOMAD: it is only through the use of nonlinear simulations that we could conduct such quality checks and glean insights into the applicability of the different designs.

## Discussion

Rational strain design using kinetic models is one of the holy grails of metabolic engineering since it obviates the need for expensive high-throughput experiments and provides a structured approach to strain design. NOMAD provides a systematic framework to achieve this by using a first-principles led approach to build quality kinetic models and then conduct rational strain design using a judicious choice of design specifications and constraints.

Although several frameworks exist to produce kinetic models that are representative of *steady-state* behavior, we demonstrate the need for carefully choosing models based on their *dynamics* as well. Through our multi-step screening process, we use fundamental engineering principles to obtain high quality kinetic models that not only reproduce the dynamics of the reference strain but also capture the trends observed during the implementation of experimental engineering strategies.

Using the provided case study, we have also shown that it is crucial to maintain the phenotype of the engineered strains close to the reference strain. Small phenotype perturbation ensures the reliability of the obtained engineered strains and provides superior titers of the desired biochemical. One way to minimize perturbations manually through MCA would be to choose combinations of enzymes that have strong control over the production pathway with negligible negative impact on growth or other system critical pathways. However, such an approach will still be unable to control the deviations in the individual concentrations, or predict the impact of perturbing multiple enzymes together. By using NRA, we circumvent these issues and provide a scalable and computationally efficient way to use the information contained in the control coefficients to constrain the network effects of the proposed changes while achieving the desired metabolic objective.

With high-quality models, it is tempting to assume that the strain design process is seamless. On the contrary, we have shown that avoiding the inherent combinatorial explosion when conducting unguided and unbiased rational strain design is not trivial. We achieve this by judiciously framing an optimization problem around control coefficients to efficiently enumerate multiple routes to achieve the design objective. We then provide a systematic way to validate these designs, highlighting the need for rigorous robustness evaluation and sensitivity analysis to ensure that the proposed designs perform well across phenotype and expression uncertainties. Lastly, the models provided in this study can also be used for strain design through nonlinear optimization.

Overall, NOMAD presents a versatile, modular framework whose concepts are applicable regardless of the size of the model, the type of kinetic mechanisms used, or the framework used to build the putative models. In doing so, it paves the way for accelerating the use of kinetic models in strain design endeavors.

## Methods

### Generating putative kinetic models of E. coli W3110 trpD9923 using ORACLE

As a case study, we use kinetic models to propose rational design strategies for the overproduction of anthranilate in a strain of *E. coli W3110 trpD9923*. This is a strain that accumulates anthranilate due to a loss of anthranilate phosphoribosyltransferase (ANPRT) activity leading to tryptophan auxotrophy. To build such kinetic models, we require knowledge of the reaction mechanisms, and the parameters that characterize each mechanism. The complexity of the metabolic network coupled with physiological and parametric uncertainty, renders this a challenging task. To overcome these challenges, we used the ORACLE framework^15,16,28,39,40^ to develop a set of putative kinetic models that represent the strain.

#### Reduced model generation

To build a kinetic model, we first need stoichiometric information about the metabolic network. We used redGEM^41^ and lumpGEM^42^ to create a reduced model of *E.coli*, and then removed the ANPRT reaction to mimic the nonsense mutation in *trpD9923*. We retained all reactions belonging to the core subsystems – glycolysis, pentose phosphate pathway (PPP), the Krebs cycle, anaplerotic reactions, the shikimate pathway, and glutamine synthesis, and added a single reaction for growth by lumping the biosynthetic reactions. The resulting network had 196 reactions (with 81 transport reactions) and 159 metabolites, spread across 2 compartments, the cytoplasm and the periplasm.

#### Data integration – metabolomics, fluxomics & thermodynamics

Before generating samples of steady-state concentrations and fluxes, we integrated exo-metabolomic and exo-fluxomic information obtained at the start of the exponential phase for the reference *W3110 trpD9223* strain^27^. Since no lag-phase was observed in this strain, this corresponded to the start of the fermentation process itself. For the glucose uptake rate, and the growth rate, we fitted analytical batch fermentation curves to the experimental data^27^. We used information on the M9 minimal medium content to constrain the extracellular metabolite concentrations. In addition to this, we integrated general metabolomics and thermodynamics data^43–45^.

#### Sampling of steady-state concentrations and fluxes

To ensure that the sampled steady-state profiles had thermodynamically consistent reaction directionalities, we used thermodynamics-based flux balance analysis (TFA)^46^ implemented in pyTFA^47^ to generate 4000 steady-state samples that resulted in at least 80% of the maximal growth. These samples consist of fluxes, concentrations, and thermodynamic variables associated with each reaction (*ΔG ‘°,ΔG*”).

#### Data integration – kinetic reaction mechanisms

Depending on the stoichiometry of each reaction in the metabolic network, we assigned a reaction mechanism (Supplementary note I). The primary mechanisms we used were the Generalized Reversible Hill^48^, and Convenience kinetics^49^, both of which capture enzyme saturation. We used mass action kinetics to model periplasm to extracellular transports.

Considering the importance of regulatory networks within the cell, we also modelled four types of allosteric regulation: (i) competitive inhibition, (ii) uncompetitive inhibition, (iii) mixed inhibition, and (iv) activation. We obtained regulatory information from an earlier study^50,51^ . We then added the regulation of DDPA and ANS by end-product metabolites. In total, we incorporated regulatory information for 31 reactions, including interactions for 5 reactions in the Shikimate pathway (Supplementary note I).

#### Kinetic model generation

With the stoichiometry and reaction mechanisms at hand, we needed to determine the kinetic parameter sets that characterize the system of ODEs using the ORACLE framework^15^. For each of the 4000 steady-state samples, we sampled 200 sets of kinetic parameters that were consistent with the concentrations and thermodynamic displacements of each reaction. Each combination of a steady-state profile and its associated kinetic parameter set constituted one kinetic model. Next, we pruned the kinetic models for linear stability – only those models whose Jacobian matrix had all negative eigenvalues were retained. In this manner, we generated 800,000 kinetic models that were linearly stable around the reference steady-states.

### Screening kinetic models

Once the initial set of kinetic models is available, we screen them to find the ones that are representative of the dynamic characteristics of the reference strain. The screening process consists of several steps, with each step enhancing the quality of the models that satisfy the requisite criteria.

Linearized dynamics: We built each kinetic model from the initial set around a steady-state consistent with the integrated experimental data. However, not all of these models necessarily capture the experimentally observed dynamics of the metabolic network. To identify models with physiologically-relevant dynamic properties, we assume that: (i) any experimentally observable steady-state is locally stable; and (ii) since metabolic reactions occur at a timescale of seconds and milliseconds, metabolic processes should settle before the cell division, which is at a timescale of minutes and hours.

To this end, we first linearize the models around their steady-states, and estimate the time constants using the eigenvalues of the Jacobian. To compute the Jacobian, we need the kinetic parameters computed by a kinetic modeling technique and the steady-state concentrations in the metabolic network. These concentrations can be obtained by integrating the set of ODEs till they reach a steady-state as done by MASSpy^17^ or Ensemble Modeling (EM)^19^, or directly from the constraint-based models used to build the kinetic models as in pyTFA^47^. We then use these calculated time constants to screen the models. Assuming aperiodic responses to perturbations, models returning to within 1% of their original steady-states by the doubling time of the cell should have dominant time constants at least five times smaller than the doubling time.

For the case study presented in this work, we chose models with a dominant time constant of less than 25 minutes.

#### Nonlinear response to concentration perturbations

The above linear stability analysis provides information on how the network will respond to infinitesimally small perturbations. However, in actual fermentation settings, the cell traverses different phases, such as the lag and exponential phases, during which there are significant fluctuations in concentration profiles. In the presence of experimental fermentation data, we can directly verify the robustness of our models to these fluctuations by checking if they can reproduce the fermentation curves. However, in the absence of such temporal data, we verify the robustness of the models using their nonlinear responses to randomly applied concentration perturbations instead. To do this, we apply a ‘k-fold’ perturbation to the steady-state concentrations of each kinetic model and integrate the system of ODEs to verify whether or not the perturbations are damped out before the cell’s doubling time. We repeat this ‘n-times’ and select those models for which all the perturbed models return to the original steady-state within the physiological timescale of the cell.

This step was unnecessary for the current study since we had ample fermentation data to compare against our results.

#### Reproduction of batch fermentation data

In this phase of model screening, we integrate information about the inoculum, and the fermentation medium, and run batch fermentation simulations using the models that are selected in the previous step. We then choose those models that can accurately capture experimental fermentation data, which is available in the form of growth curves, secretions, and uptakes.

For obtaining kinetic models representative of *E.coli W3110 trpD9923*, we integrated inoculum information provided in the study^27^ and ran batch fermentation simulations for each of the screened models. We then chose those models within 5% and 10% of the final steady-state values of growth and extracellular anthranilate, respectively, and whose fermentation times were less than 20 hours.

#### Robustness to enzymatic interventions

The end goal of the framework is to provide targets for enzymatic interventions that enable us to achieve a given metabolic output. Not all the models that are selected in the previous screening steps are equally robust to enzymatic interventions - some can veer significantly from their behavior, showing little to no growth, while others can retain their reference growth level. Hence, to determine the robustness of each kinetic model to such interventions we apply a ‘k-fold’ perturbation to the maximal velocities of the different reactions and study the growth of the resulting strain. We repeat this ‘n’ times and choose those models for which all the perturbed strains demonstrate satisfactory growth.

In the presented case study, we applied a 10% normally distributed perturbation to the maximal velocities of each reaction in the network and then integrated the system of ODEs in a batch reactor setting. We repeated this process 50 times for each kinetic model and chose those models that displayed at least 50% of the experimentally observed biomass for all the 50 perturbations.

The end product of this 4-step filtering process is a *population* of robust, representative kinetic models that are adequate for rational strain design.

### Robust strain design using kinetic models

We use the screened kinetic models to conduct rational strain design with a given objective to be attained. For the presented case study, the objective was to maximize the yield of anthranilate with respect to glucose uptake. The strain design process can be divided into the following steps:

#### Generating design alternatives using Network Response Analysis

One approach to strain design would be to exhaustively simulate all possible combinations of target enzymes along with the degrees of up or down regulations applied to them. The arduous nature of this task and the computational cost involved provide a strong case for a more judicious approach to choosing enzymatic targets.

One possibility would be to use Metabolic Control Analysis (MCA)^34,35,52^, i.e., to calculate the log-linear sensitivities of the production pathway to system parameters and to then use the enzymes with the top control coefficients as the candidates to be tested in a nonlinear setting. However, this approach has its drawbacks. An increase in enzyme activity affects not only the target flux/metabolite but also other components of the network, potentially causing a significant deviation from the reference physiology, or the accumulation of toxic metabolites. This situation is further complicated when targeting multiple enzymes simultaneously. A starting point to overcome such deleterious effects would be to use heuristics and expert knowledge and eliminate from contention those targets that are known to have undesirable network effects.

To provide a more systematic and efficient approach to dictate such choices and constraints, a constraint-based MCA method called Network Response Analysis (NRA) was developed^26^. In NRA, we frame the strain design objective as a mixed-integer linear optimization problem built around the control coefficients, and the reference steady-state profiles of concentrations and fluxes. In addition, we supply design constraints such as the allowable fold-change in fluxes, concentrations, and enzyme activities, and the number of allowable enzymatic interventions. In this way, NRA provides two distinct advantages. First, it ensures the reliability and robustness of designs by controlling the deviation from the reference phenotype through the imposed constraints. Second, by using an optimization problem, NRA provides a computationally efficient and scalable approach to strain design by avoiding the combinatorial explosion inherent when we seek multiple enzymatic targets.

With these features in mind, we use NRA to enumerate designs for each of the chosen kinetic models that achieve the desired objective within a certain threshold. In this manner, we can generate hundreds of designs across all the kinetic models.

In the current study, we imposed the following constraints: (i) a maximum of 3-fold change in concentrations and 5-fold change in enzyme activities, (ii) a maximum of 3 enzymatic interventions, and (iii) a maximum of 20% reduction in growth rate. We set the objective to be the maximization of anthranilate yield with respect to glucose uptake and enumerated all designs within 5% of the maximal objective for each kinetic model.

#### Design ranking

At the end of previous step, we have a list of putative designs generated using each kinetic model. As we have a population of kinetic models as opposed to a single representative model, we need to carefully choose those designs that are robust across the entire population. Robustness can be characterized in many ways – the reappearance of the same design by membership across different models, or the highest predicted objective when the design is enforced across different models etc. The criteria to define robustness can vary with the objective that we seek to attain.

In this work, we first extracted those designs that were unique by membership and enforced them in each of the kinetic models by setting the minimal log fold change in the enzyme activity levels of the enzymatic targets to be 1e-6. We then ranked the designs by the mean predicted increase in anthranilate yield with respect to glucose across all models, and selected the top 5 designs as the most robust strategies.

#### In silico design verification and sensitivity analysis

Once we have ranked and chosen the most robust designs, we verify them *in silico* in a batch fermentation setup and study the sensitivity of the designs to vagaries in experimental implementation. This step is necessary since the performance of the designs in the previous step was evaluated based on a log-linear approximation of the system, taken at the reference steady-state. By verifying the proposed designs in a nonlinear setup, we can understand how well the log-linear approximations fare in a nonlinear setting. After the design verification and analysis, the most promising designs are sent for experimental implementation and validation.

For the presented case study, we first analyzed the performance of the top 5 designs in a batch fermentation setting, using the NRA predicted enzyme fold changes specific to each model. For the inoculum and medium, we integrated the same information as was done in the model screening step. For the sensitivity analysis, we first calculated the mean NRA-suggested fold changes for each enzyme for a given design. We then applied a ±50% uniformly distributed perturbation to the mean fold changes for (i) all 3 enzymes, (ii) each enzyme individually while keeping the other two enzymes at the mean NRA-suggested fold changes. To obtain a statistical estimate of the sensitivities, we did this 10 times for each of the 10 models and tracked the mean across the 100 responses for each design.

## Supporting information

Reaction information

## Acknowledgements

This work was supported by funding from the Swiss National Science Foundation Synergia grant CRSII5_198543, the European Union’s Horizon 2020 research and innovation programme under grant agreement 814408, Swedish Research Council Vetenskapsradet grant 2016-06160, and the Ecole Polytechnique Fédérale de Lausanne (EPFL).

