## Supplementary notes for "Rational strain design with minimal phenotype perturbation"

### Supplementary Information

#### Supplementary note I – reaction information

Information about the rate laws and regulatory information used for each reaction are provided in the supplementary file, reaction_information.xlsx. The inhibitions were modelled as follows for a sample Michaelis-Menten equation. Further information on the way reaction mechanisms are modelled can be found in the supplementary information section of SKiMPy [1].

a) Simple inhibition:

$$v = V_{max} \cdot \left(1 - \frac{\Gamma}{K_{eq}}\right) \cdot \left(\frac{\frac{S}{K_S^M}}{\frac{S}{K_S^M} + \frac{P}{K_P^M} + 1}\right) \cdot \left(\frac{\mathbf{1}}{\mathbf{1} + \frac{\boldsymbol{I}}{\boldsymbol{K_I}}}\right)$$

b) Competitive inhibition:

$$v = V_{max} \cdot \left(1 - \frac{\Gamma}{K_{eq}}\right) \cdot \left(\frac{\frac{S}{K_S^M}}{\frac{S}{K_S^M} + \frac{P}{K_P^M} + 1 + \frac{\boldsymbol{I}}{\boldsymbol{K_I}}}\right)$$

c) Mixed inhibition:

$$v = V_{max} \cdot \left(1 - \frac{\Gamma}{K_{eq}}\right) \cdot \left(\frac{\frac{S}{K_S^M}}{\frac{S}{K_S^M} + \frac{P}{K_P^M} + 1 + \frac{\boldsymbol{I}}{\boldsymbol{K_I}}}\right) \cdot \left(\frac{\mathbf{1}}{\mathbf{1} + \frac{\boldsymbol{I}}{\boldsymbol{K_I}}}\right)$$

d) Simple activation:

$$v = V_{max} \cdot \left(1 - \frac{\Gamma}{K_{eq}}\right) \cdot \left(\frac{\frac{S}{K_S^M}}{\frac{S}{K_S^M} + \frac{P}{K_P^M} + 1}\right) \cdot \left(\mathbf{1} + \frac{\boldsymbol{A}}{\boldsymbol{K_A}}\right)$$

#### Supplementary note II – top 5 designs

The mean NRA proposed fold changes in enzyme activity for the top 5 designs, when each of them was enforced for the 10 kinetic models are given in the table below. The designs all propose strongly upregulating DDPA, while reducing the activity of GLUDy by around 20%. For all the designs, we used the usual constraints – maximum 5-fold upregulation, unlimited downregulation, maximum 3-fold concentration change and maximum 3-enzyme modifications.

| Design | Enzyme | Mean suggested fold-change across 10 kinetic models |
|---|---|---|
| **d-1** | DDPA↑ | 4.15 |
| | GLUDy↓ | 1.2 |
| | PYK↑ | 1.57 |
| **d-2** | DDPA↑ | 3.91 |
| | GLUDy↓ | 1.19 |
| | PGI↓ | 2.23 |
| **d-3** | DDPA↑ | 3.92 |
| | GLUDy↓ | 1.2 |
| | GND↓ | 1.62 |
| **d-4** | DDPA↑ | 3.89 |
| | GLUDy↓ | 1.21 |
| | HEX1↓ | 3.28 |
| **d-5** | DDPA↑ | 2.48 |
| | GLUDy↓ | 1.13 |
| | ANS↑ | 2.87 |

#### Supplementary note III – addendums to Figure 7 (main text)

Four out of the top 5 designs, d-1 to d-4, proved to be robust across the 10 kinetic models when implemented using the NRA suggested fold changes in enzyme activity specific to each model. To ensure their suitability for experimental implementation, we tested the response of these four designs to errors in enzyme expression levels (Figure 7 in the main text). Here we provide the glucose and biomass curves for the perturbation experiments.

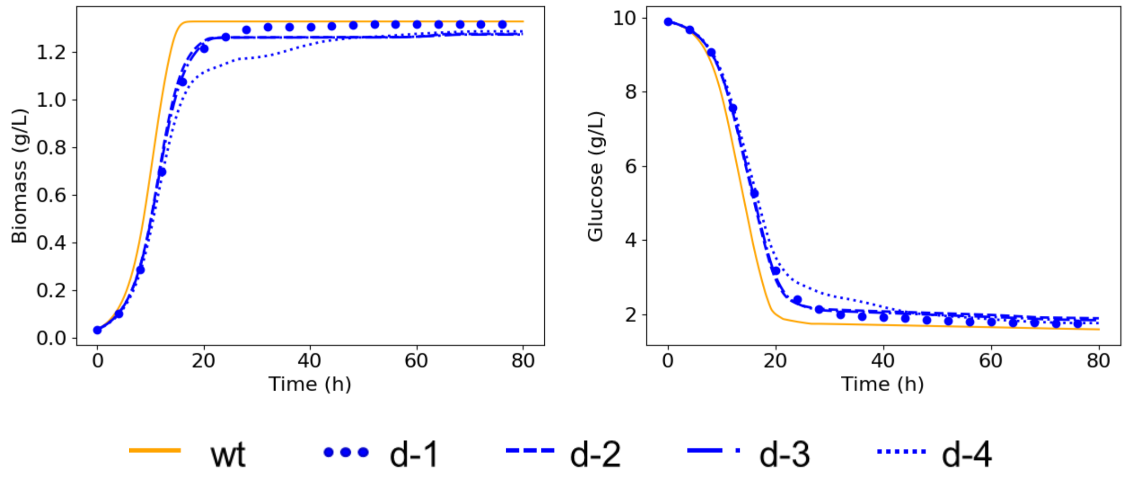


*Figure S.1: Mean biomass and glucose curves when we perturb all 3 enzymes simultaneously by* $\pm 50\%$ *(main text).*

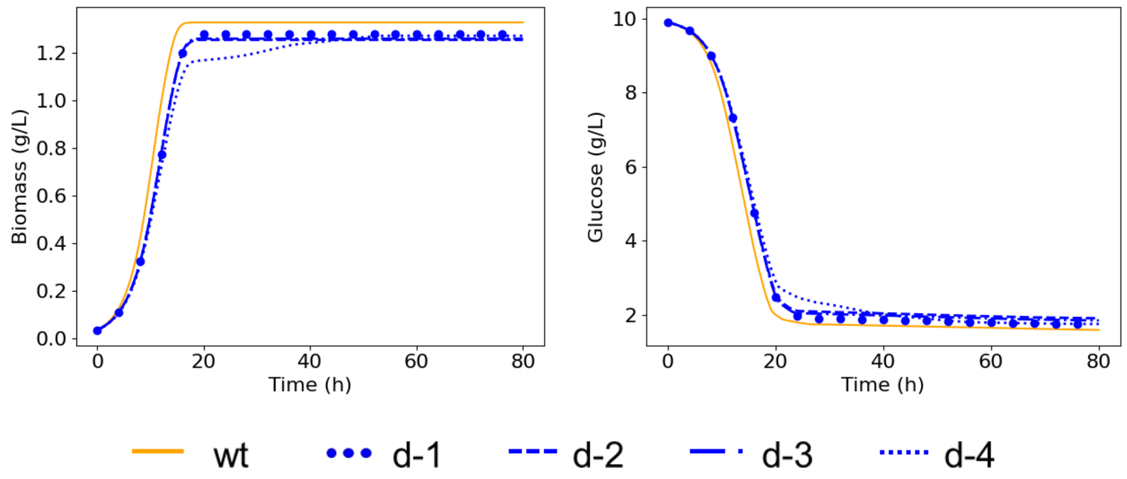


*Figure S.2: Mean biomass and glucose curves when we perturb DDPA alone by* $\pm 50\%$.

Figure S.1 shows the mean glucose and biomass curves for each of the four designs across the 10 models when we perturb all three enzymes by $\pm 50\%$, 10 times for each model. Figure S.2 shows the responses when we perturb DDPA upregulation alone while fixing the GLUDy regulation and the regulation of the third enzyme (PYK/HEX1/GND/PGI) at the mean NRA proposed values.

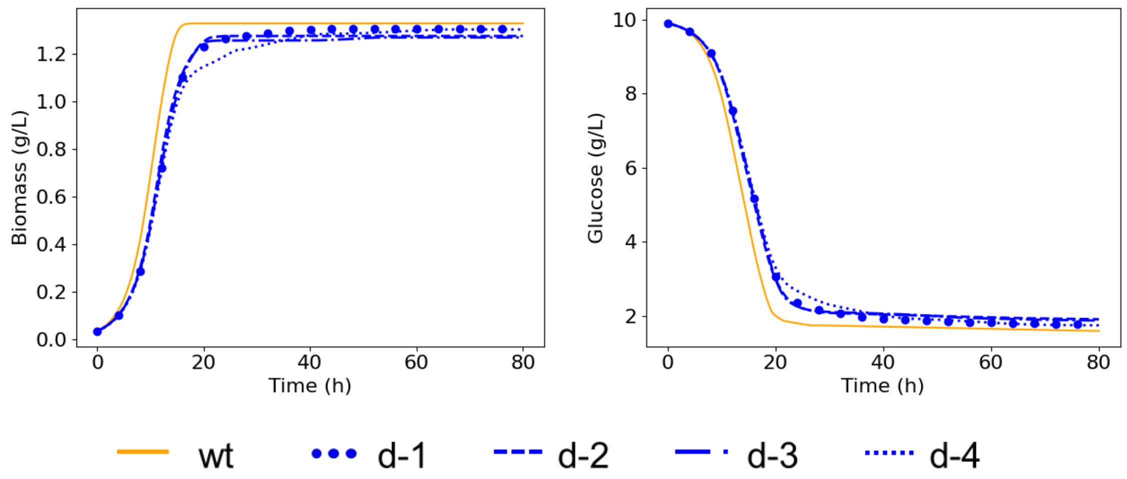


*Figure S.3: Mean biomass and glucose curves for the sensitivity analysis for GLUDy.*

Similarly, Figures S.3 and S.4 show the mean biomass and glucose curves when GLUDy and the third enzyme (either PYK, PGI, GND, or HEX1) are perturbed respectively, with the other enzymes being fixed at the mean NRA proposed fold changes. In all the perturbation experiments, we observe a negligible change in the dynamics of growth or glucose uptake, suggesting that all the four designs are suitable for experimental validation.

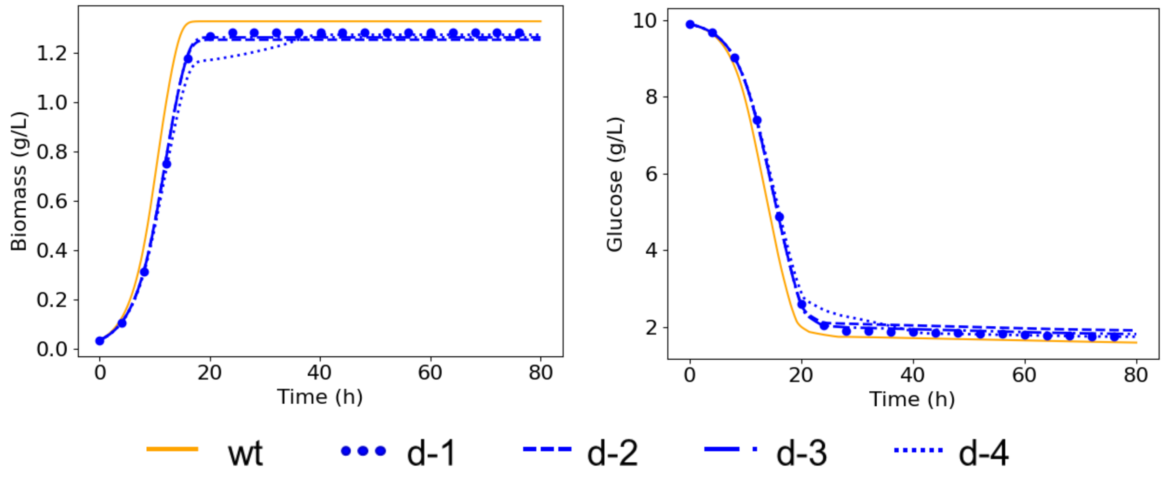


*Figure S.4: Mean biomass and glucose curves for the sensitivity analysis for the third enzyme (either of PYK, PGI, GND, or HEX1).*

### Supplementary note IV – addendums to Figure 5 (main text)

The growth and glucose curves for the simulated experimental designs and the NOMAD proposed designs are shown in Figure S.5.

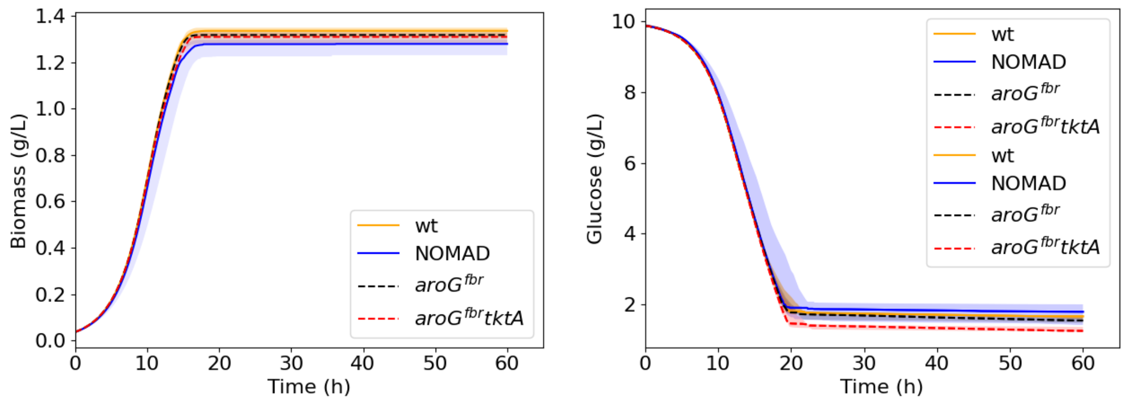


Figure S.5: Mean biomass and glucose curves across the 10 models when we implement the experimental strains, and when we implement the 67 NRA proposed designs in the specific models that were used to devise them. We see that the nonlinear simulations all follow the dynamics of the reference strain, with the only major change being the increase in anthranilate titers (main text).

#### Supplementary note V – MCA design

In the main text, we demonstrated the drawbacks of conducting strain design using the top yield control coefficients (CCs) for anthranilate with respect to glucose uptake without imposing any constraints on the phenotype perturbation. In practice, it is likely that an engineer would use their expert knowledge to judiciously choose those control coefficients that do not have an adverse impact on the network. To understand the impact of such an approach, we chose the top 3 yield CCs for each kinetic model that did not reduce growth rate. We did this by screening for those enzymes that had the same qualitative control (positive or negative) over both anthranilate and biomass yield. We then selected the top 3 enzymes among the filtered enzymes based on their anthranilate yield CC and then applied them using a 5-fold perturbation. We observed that even with this choice of targets, both growth and anthranilate production were severely hindered (Figure S.6). Indeed, it is only when we limit the allowable fold change in perturbations of enzyme activity to less than 1.5 fold (Figures S.6-8) that the designs start yielding superior anthranilate titers. This further supports NRA as a holistic, systematic approach to applying design constraints.

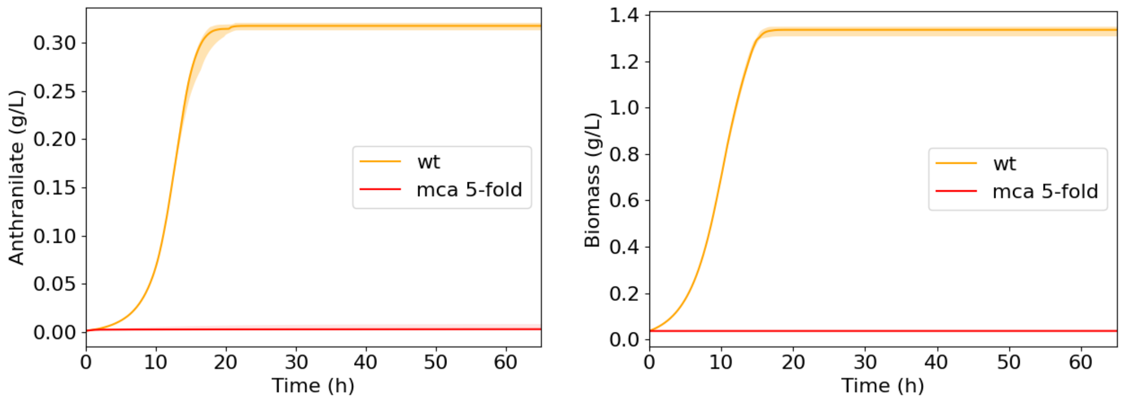


Figure S.6: Mean anthranilate and biomass curves across the 10 models for the wild type (wt) strains and MCA-designed strains when we apply a 5-fold change in enzyme activities to the top 3 anthranilate yield control coefficients that do not have an adverse effect on the growth rate. We see that both growth and anthranilate production are adversely impacted even when we account for the impact on growth rate.

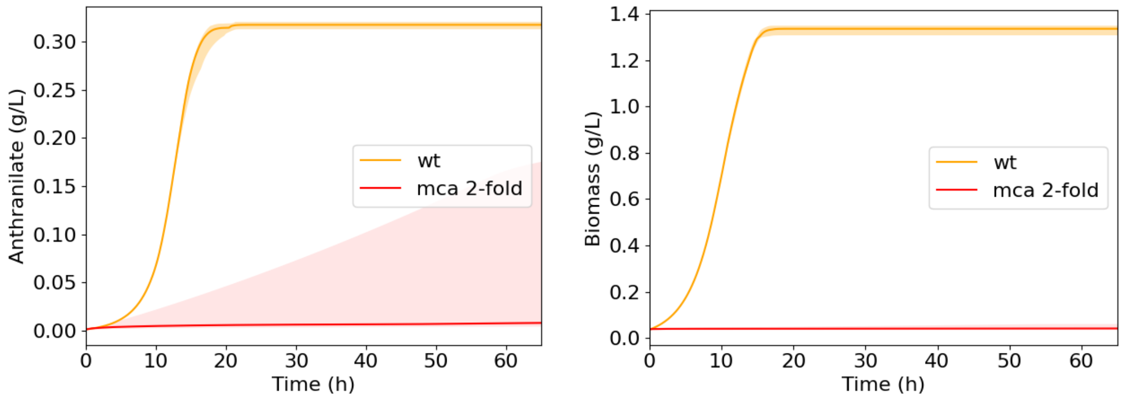


Figure S.7: Mean anthranilate and biomass curves across the 10 models for the wild type (wt) strains and MCA-designed strains when we apply a 2-fold change in enzyme activities to the top 3 anthranilate yield control coefficients that do not have an adverse effect on the growth rate. When compared with the MCA designs with 5-fold changes in enzyme activities, we see marginal improvement in anthranilate titers, although growth is still severely impacted.

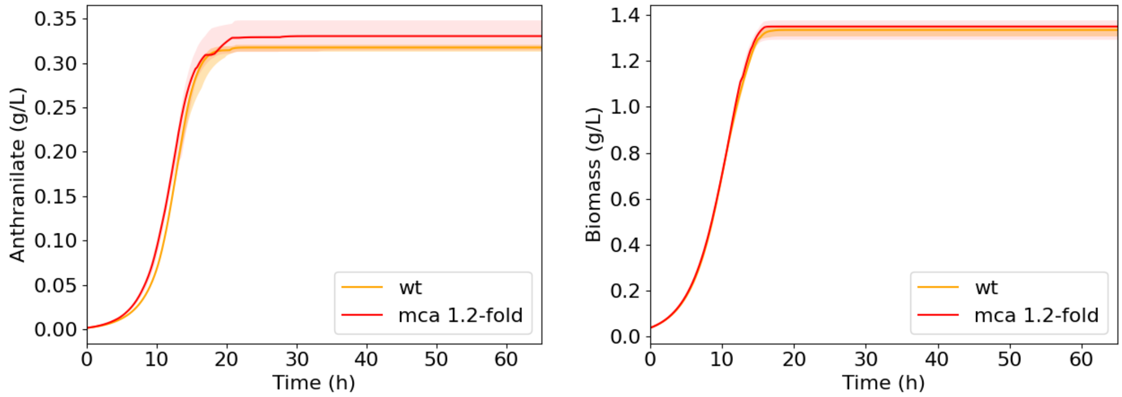


Figure S.8: Mean anthranilate and biomass curves across the 10 models for the wild type (wt) strains and MCA-designed strains when we apply a 1.2-fold change in enzyme activities to the top 3 anthranilate yield control coefficients that do not have an adverse effect on the growth rate. We see that as we constrain the allowable change in expression levels, a constrained MCA approach based on a judicious manual choice of control coefficients becomes more viable.
